# Pathology in resected areas of [^18^F]FDG PET hypometabolism in pediatric epilepsy patients with focal cortical dysplasia type II

**DOI:** 10.64898/2026.06.05.729979

**Authors:** Jack Lam, Nicolás von Ellenrieder, Meriem Hamel, Yunzhu Ruan, David Dufresne, Marie-Christine Guiot, Jason Karamchandani, Boris C. Bernhardt, Roy Dudley

**Affiliations:** Department of Neurology and Neurosurgery, Montreal Neurological Institute and Hospital, McGill University, Montreal, QC, Canada; Department of Pediatrics, Université de Sherbrooke, Sherbrooke, QC, Canada; Department of Pathology, McGill University Health Centre, McGill University, Montreal, QC, Canada; Department of Pediatric Neurosurgery, Montreal Children’s Hospital, McGill University, Montreal, QC, Canada

**Author notes:** Corresponding Author: Dr. Roy DudleyDepartment of Pediatric Neurosurgery, Montreal Children’s Hospital McGill University, 1001 Decarie Blvd. Montreal, QC H4A 3J.

**Keywords:** epilepsy, pediatric, FDG PET, pathology

## Abstract

**Introduction:** [^18^F]fluorodeoxyglucose positron emission tomography (FDG-PET) frequently reveals hypometabolism extending beyond MRI-visible focal cortical dysplasia (FCD). Whether surrounding hypometabolic areas harbour pathological cells remains unclear. We compared histopathology between the lesion epicentre and surrounding hypometabolic borders in children undergoing epilepsy surgery.

**Methods:** Fourteen children with intractable, extra-temporal focal epilepsy (mean age 9.0±5.0 years; 9 female) were retrospectively reviewed. Resections encompassed the visually-apparent MRI signal abnormalities as well as areas of surrounding FDG-PET hypometabolism when safely feasible. Multiple pathological specimens were obtained from the epicentre and surrounding hypometabolic areas. Overall, 136 specimens were analyzed: 64 epicentre (median 4±3/patient) and 72 border (median 5±4/patient).

**Results:** Pathology was identified in 75% of epicentre specimens (59% with frank FCD (fFCD) IIa/b, 16% with dysmorphic neurons only (DNO)). Border specimens showed pathology in 62% (31% fFCD IIa/b, 31% DNO). We fitted a Bayesian logistic mixed model with pathology as outcome variable, location as predictor, and subject as a random effect. Compared to negative pathology, the log-odds of fFCD in the epicentre was 1.00 (credible interval (CI) 0.32, 1.77) and −1.25 in the border (CI −2.17, −0.40). The log-odds of DNO vs negative pathology was non-significant in both locations. Exploratory analyses did not show significant associations between pathology and degree of regional hypometabolism or volume. All patients achieved Engel Ia status at one-year follow-up with no long-term neurological deficits.

**Conclusion:** These findings suggest a spatial difference in distribution of histopathology, with fFCD concentrated in the epicentre and DNO present in both the epicentre and hypometabolic borders. Thus, FDG-PET may be used to better detect the histopathological borders of FCD type II, and the high seizure-freedom rate presented here supports the inclusion of these surrounding hypometabolic regions in the surgical resection (when safe to do so), improving the likelihood of removing potentially epileptogenic cells.

**Key Points:**

1. Pathological cells are present not only in the MRI signal abnormality in FCD but also in the periphery of the FDG-PET hypometabolism.
2. We observe a spatial difference in distribution of histopathology, with frank FCD concentrated in the epicentre and dysmorphic neurons spread throughout the area of hypometabolism.
3. Maximal safe resection of the area of hypometabolism may increase likelihood of removing potentially epileptogenic cells, thus improving surgical outcome.

## Introduction

Focal cortical dysplasia (FCD) type II is the cause of seizures in 17-31% of children undergoing surgery for pharmacoresistant epilepsy.^1^ On histopathology, it is characterised by disruption of cortical architecture and dysmorphic neurons, either without (type IIa) or with balloon cells (type IIb).^2,3^ Presurgical imaging is essential for delineation of the epileptogenic lesion for planning surgical resection, which can be curative in approximately 50% of patients.^4^ Magnetic resonance imaging (MRI) often shows cortical thickening, blurring of the grey-white junction, hyperintense signal on T2/fluid-attenuated inversion recovery (FLAIR) sequence, and presence of a “transmantle sign.”^5^ However, despite advances in imaging technology, subtle FCD lesions may be missed by conventional 3 Tesla (3T) MRI, necessitating further imaging investigations.

One commonly used adjunct imaging technique in the work up of pediatric epilepsy is interictal [^18^F]fluorodeoxyglucose positron emission tomography (FDG-PET), which can enable detection of a lesion even in cases where 3T MRI investigations are ambiguous, or where its borders are difficult to demarcate ^6–8^ FDG-PET often shows hypometabolism in the epileptogenic zone (EZ) and, in one survey, provides additional information in 77% of pediatric cases and had a minor or major impact on clinical decision making in 51%.^9^ A recent systematic review and meta-analysis demonstrated that localization of the FDG-PET hypometabolism was associated with improved post-surgical outcomes regardless of presence of a lesion visible on MRI.^10^ Furthermore, the extent of FDG-PET abnormalities had prognostic value as well, with diffuse hypometabolism having worse surgical outcome compared to focal hypometabolism.

While sensitive to the EZ, it is well established that the spatial extent of the FDG-PET hypometabolic abnormality may extend beyond any visible lesion on 3T MRI.^11–13^ The pathophysiology underlying this hypometabolism (i.e., underutilization of glucose) in epilepsy is poorly understood. There is emerging evidence that epilepsy is associated with abnormalities in cellular and mitochondrial metabolism with bidirectional links between metabolism and seizures.^14^ Chronic epileptic tissue shows impaired ATP production capacity,^15^ which may be secondary to impairment of proteins involved in mitochondrial function, such as complex IV.^16^

It is currently unclear whether areas of hypometabolism beyond the MRI-visible lesion reflects simply a functional deficit zone or if it might also harbour pathological cells which could contribute to epileptogenicity. In this study of pediatric epilepsy patients whose surgical plans included peripheral areas of hypometabolism, we sought to assess the presence and degree of pathology in the epicentre of the EZ as well as the borders of the hypometabolism-informed resection.

## Methods

### Patients

This study was approved by the research ethics board (REB) of the McGill University Health Centre. All patients and/or their parents signed an informed consent form for participation in the study. We reviewed 48 consecutive pediatric patients who underwent resective surgery for epilepsy by a single surgeon (R.W.R.D.) at the Montreal Children’s Hospital between 2015-2024. We included patients whose surgical plans were based on resecting 3T MRI signal abnormalities as well as surrounding hypometabolism on FDG-PET, whose histopathology was consistent with FCD, and who had greater than 1 year post-surgical follow up. All patients in this study had signal abnormalities on 3T MRI consistent with FCD, but in some cases this was only seen in retrospect upon re-examining these MRIs in the context of other presurgical evaluation findings (e.g., FDG-PET, MEG) and/or intracranial recordings. Patients with negative histopathological findings were excluded from the study. Furthermore, patients with mesial temporal lesions were excluded as it was difficult to define the borders of the lesion.

In the end, 14 pediatric patients (mean age 9.0±5.0 years, 9 females) were included in the study. Clinical characteristics of the patient sample can be found in **Table 1**. All patients were admitted for extensive presurgical work up including 3T MRI, FDG-PET, video electroencephalography (vEEG), semiology analysis, and neuropsychology. The majority of patients also underwent further investigation with interictal/ictal single photon emission computed tomography (SPECT) and magnetoencephalography (MEG). Two patients underwent stereo-electroencephalogaphy (SEEG) explorations as clinically indicated, while one patient underwent subdural grids and depth electrodes. All cases were discussed at the multidisciplinary seizure conference and resective surgery was recommended if a strong consensus for a putative EZ was reached.

**Table 1.**
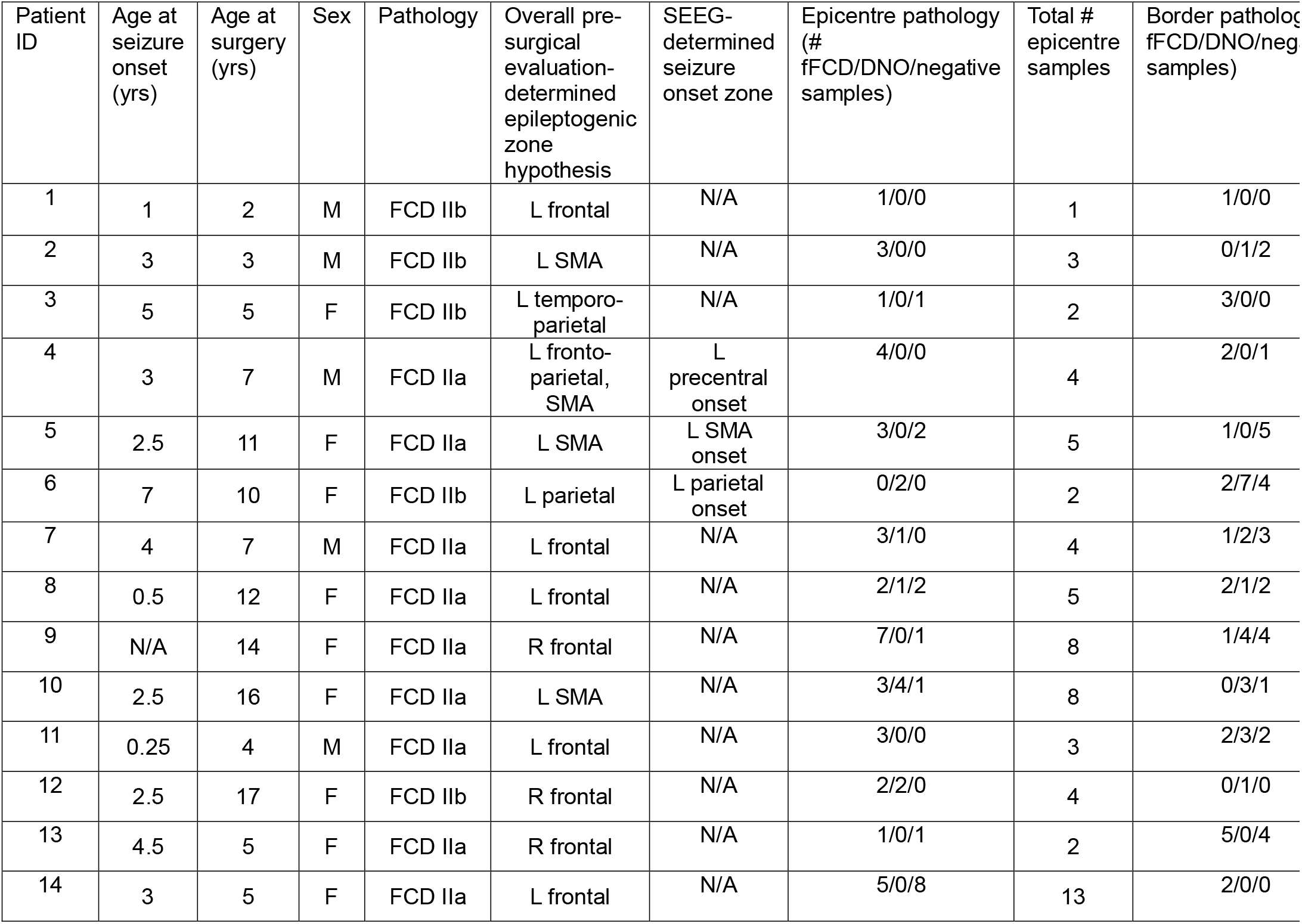
Patient characteristics and summary of pathology findings. DNO, dysmorphic neurons only; F, female; fFCD, frank focal cortical dysplasia; ID, identification; L, left; M, male; N/A, not available; R, right; SMA, supplemental motor area; yrs, years.

| Patient ID | Age at seizure onset (yrs) | Age at surgery (yrs) | Sex | Pathology | Overall pre-surgical evaluation-determined epileptogenic zone hypothesis | SEEG-determined seizure onset zone | Epicentre pathology (# fFCD/DNO/negative samples) | Total # epicentre samples | Border patholog[illegible] fFCD/DNO/neg[illegible] samples) |
|---|---|---|---|---|---|---|---|---|---|
| 1 | 1 | 2 | M | FCD IIb | L frontal | N/A | 1/0/0 | 1 | 1/0/0 |
| 2 | 3 | 3 | M | FCD IIb | L SMA | N/A | 3/0/0 | 3 | 0/1/2 |
| 3 | 5 | 5 | F | FCD IIb | L temporo-parietal | N/A | 1/0/1 | 2 | 3/0/0 |
| 4 | 3 | 7 | M | FCD IIa | L fronto-parietal, SMA | L precentral onset | 4/0/0 | 4 | 2/0/1 |
| 5 | 2.5 | 11 | F | FCD IIa | L SMA | L SMA onset | 3/0/2 | 5 | 1/0/5 |
| 6 | 7 | 10 | F | FCD IIb | L parietal | L parietal onset | 0/2/0 | 2 | 2/7/4 |
| 7 | 4 | 7 | M | FCD IIa | L frontal | N/A | 3/1/0 | 4 | 1/2/3 |
| 8 | 0.5 | 12 | F | FCD IIa | L frontal | N/A | 2/1/2 | 5 | 2/1/2 |
| 9 | N/A | 14 | F | FCD IIa | R frontal | N/A | 7/0/1 | 8 | 1/4/4 |
| 10 | 2.5 | 16 | F | FCD IIa | L SMA | N/A | 3/4/1 | 8 | 0/3/1 |
| 11 | 0.25 | 4 | M | FCD IIa | L frontal | N/A | 3/0/0 | 3 | 2/3/2 |
| 12 | 2.5 | 17 | F | FCD IIb | R frontal | N/A | 2/2/0 | 4 | 0/1/0 |
| 13 | 4.5 | 5 | F | FCD IIa | R frontal | N/A | 1/0/1 | 2 | 5/0/4 |
| 14 | 3 | 5 | F | FCD IIa | L frontal | N/A | 5/0/8 | 13 | 2/0/0 |

### FDG-PET and Surgical Planning

FDG-PET was obtained in systematic fashion for all patients as part of our standard focal epilepsy presurgical evaluation protocol.^17^ Seven patients at the Centre hospitalier universitaire Sainte-Justine and five patients at the Centre hospitalier universitaire de Sherbrooke were scanned on a Philips Gemini 16 time-of-flight PET/CT scanner (Amsterdam, Netherlands) with a resolution of 4.7mm. Two patients at the Centre hospitalier universitaire de Sherbrooke were scanned on a GE Discovery MI scanner (Boston, USA) with a resolution of 4.8mm. All patients fasted for 4 hours prior to the scan. Blood glucose was measured prior to [^18^F]FDG injection to ensure that levels were ≤8mmol/L. Patients were injected with a 3.5-5MBq/kg dose of [^18^F]FDG followed by a 45-60 minute acquisition. Scans from the Philips Gemini 16 time-of-flight PET/CT scanner were reconstructed using a line-of-response row-action maximum likelihood (LOR-RAMLA) algorithm, while scans from the GE Discovery MI scanner were reconstructed using Q.clear algorithm.

FDG-PET images were imported to BrainLab (Munich, Germany) software where it was co-registered to the preoperative 3T MRI.

For surgical planning, two virtual objects were manually created (i.e., segmented) by the lead surgeon (R.W.R.D) using the BrainLab software, designating (1) the visible MRI signal abnormalities (i.e., the epicentre) as well as (2) the surrounding FDG-PET hypometabolism borders, staying away from any perceived eloquent brain areas (e.g., language areas or motor areas of the precentral gyrus). **Figure 1** shows a representative patient with a left central operculum FCD with a T1 hypointense (**Figure 1A**) and FLAIR hyperintense (**Figure 1B**) lesion outlined in red. The area of FDG-PET hypometabolism is outlined in white (**Figure 1C**).

**Figure 1.**
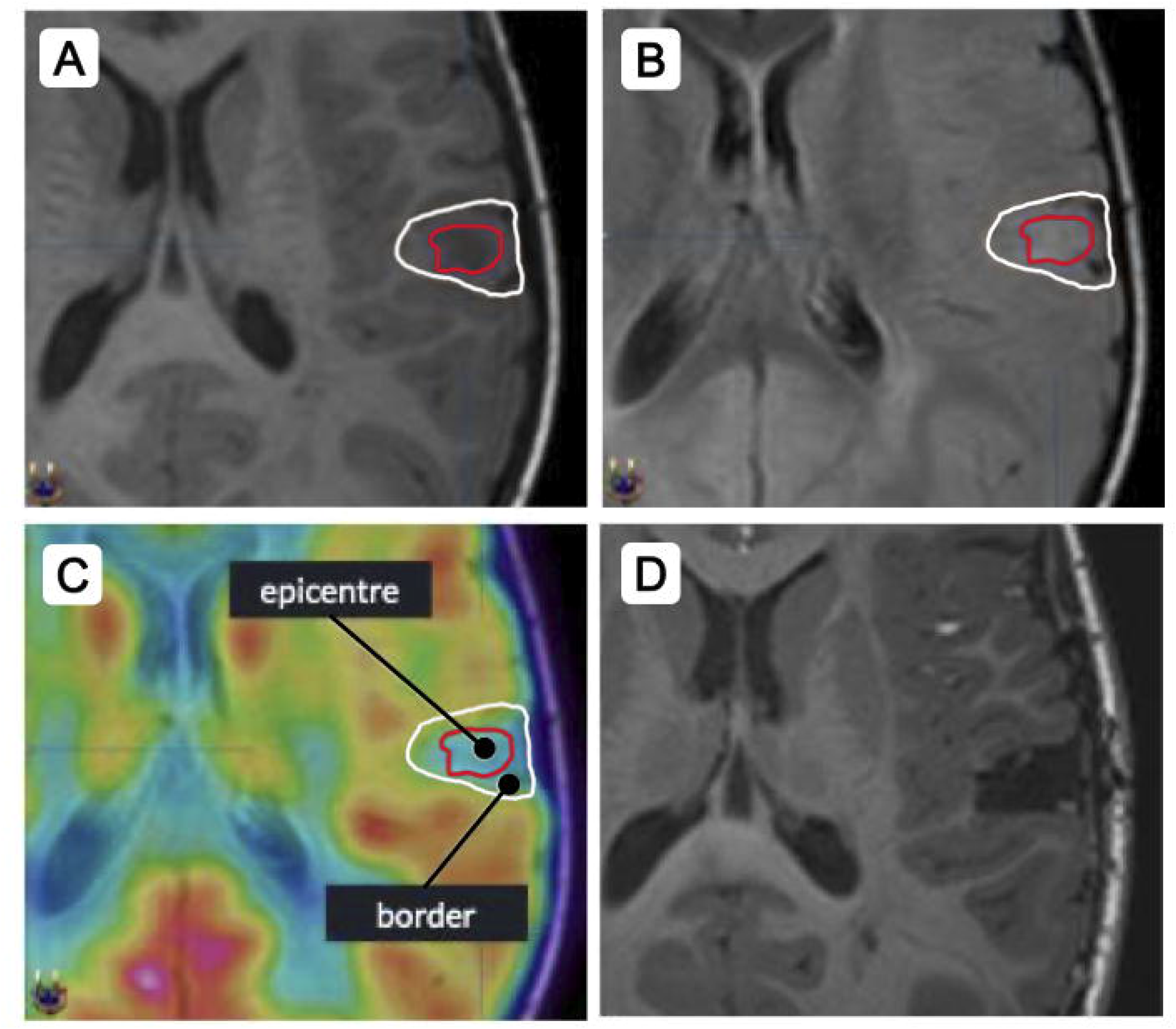
Representative case (Patient 1) with a left central operculum focal cortical dysplasia (FCD) that is hypointense on T1 (A) and hyperintense on FLAIR (B). The red line delineates MRI-visible lesion while the white line delineates the area of hypometabolism as seen on [^18^F]fluorodeoxyglucose positron emission tomography (FDG-PET) (C). Numerous intraoperative samples were obtained from each patient within the red line (Epicentre samples) and between the red and white lines (Border samples). Intra-operative MRI was done to ensure complete resection of the FDG-PET hypometabolism (D).

### Intraoperative

The area of hypometabolism served as the borders of the resection in all included patients when safely feasible. The goal of each resection was to remove both the epicentre and the borders. Intraoperatively, once the head was positioned and fixed in pins of the Noras head coil, a predissection intraoperative MRI (iMRI) was performed and coregistered to the pre-operative MRI containing the virtual objects. Then, automatic registration was performed using the fiducials contained in the Noras Head coil so that highly accurate registration was obtained for precise neuro-navigation allowing us to find the MRI epicentre of the lesion and the FDG-PET hypometabolism borders via the virtual objects. During the surgery, numerous pathological samples were taken from both the epicentre of the putative EZ (i.e., the area showing MRI signal abnormalities), and the border areas of hypometabolism (i.e., the area outside of the epicentre but within the area of hypometabolism as delineated on BrainLab). An intradissection iMRI was used to ensure that the entirety of the epicentre and surrounding hypometabolic area was resected (**Figure 1D**). If MRI abnormalities or FDG-PET hypometabolism remained according to the virtual objects, further surgery was performed immediately in the same setting, and another iMRI was performed to confirm complete resection.

### Pathology

All samples were sent for histopathological analysis by the board-certified neuropathologists (M.-C.G., J.K.). Each sample was classified as either i) frank FCD (fFCD, either type IIa or IIb) meaning that disruption of cortical architecture was seen, as well as abundant abnormal cells (dysmorphic neurons +/- balloon cells), ii) dysmorphic neurons only (DNO; presence of dysmorphic neurons (usually less abundant than what is seen in fFCD) without a clear disruption of cortical architecture to classify it as FCD and without balloon cells), or iii) negative for pathology (Neg). Samples containing white matter only or ultrasonic aspirator contents were excluded.

### Imaging Analysis

The MRI-visible lesion (epicentre) was manually segmented on Freeview software based on the abnormal signal seen on T1w and FLAIR. The postoperative T1w MRI was linearly co-registered to the preoperative T1w MRI using Label-Augmented Modality-Agnostic Registration (LAMAReg).^18^ The resection cavity was automatically segmented using MELD-PostOp^19^ and manually corrected, taking into account postoperative brain shift. All segmentations were reviewed for accuracy by the lead surgeon (R.W.R.D).

Cortical surface reconstruction and anatomical segmentation of preoperative T1w MRI was performed using FastSurfer v2.0.0.^20,21^ FDG-PET images were linearly co-registered to T1w MRIs using FreeSurfer mri_coreg. All segmentations and registrations were visually inspected for accuracy. Partial volume correction was performed using the Müller-Gärtner technique as implemented in the PETPVC toolbox.^22^ An eroded pons region-of-interest was used as the reference region for image intensity normalization. We projected the partial volume corrected standard uptake value ratio (SUVR) images, the epicentre lesion label, and the resection cavity label to mid-cortical thickness fsLR32k surface space using Connectome Workbench.^23^ FDG-PET data were smoothed with a 5mm full-width-at-half-maximum (FWHM) Gaussian kernel along the surface. The average SUVR in the epicentre and the border (the resection cavity label excluding the epicentre) was calculated on the surface. Surface asymmetry index maps were subsequently calculated using the following formula:

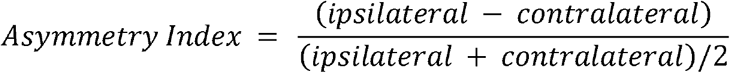

Asymmetry index maps were thresholded at < −0.20 and clustered. We designated clusters with any overlap with the resection cavity as resection-associated hypometabolic clusters. Percentage of hypometabolism resected was calculated by dividing the surface area of hypometabolic clusters within the resection mask by the total surface area of resection-associated hypometabolic clusters. In **Figure 2** we show imaging findings and surface hypometabolic clusters from three example patients: one with a subtle lesion that was seen retrospectively after intracranial EEG (Patient 5), one with a large hypometabolic abnormality (Patient 8), and one with a focal hypometabolic abnormality (Patient 14).

**Figure 2.**
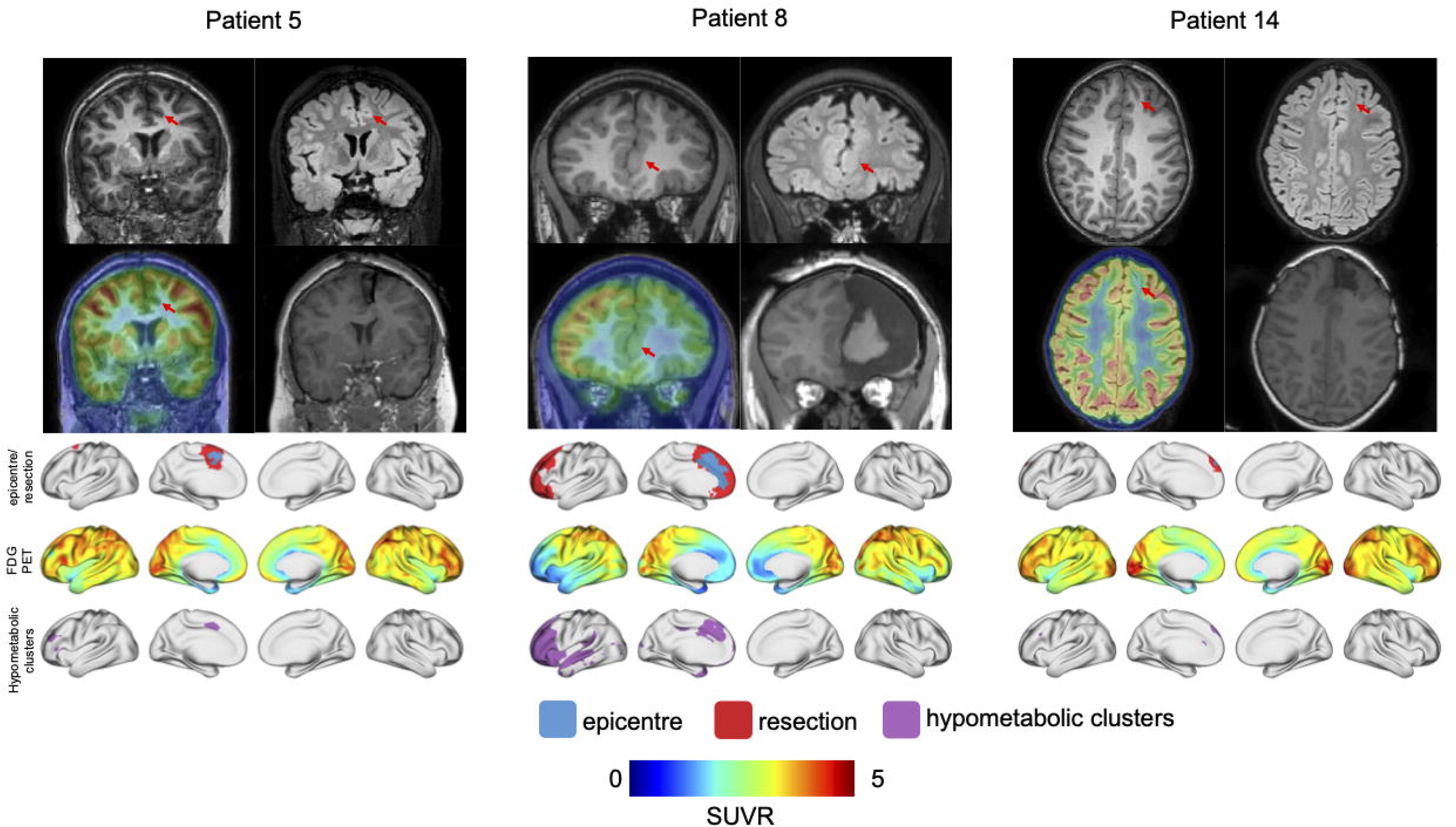
Example cases showing a left supplementary motor area, a large left frontal, and a small left frontal lesion (Patients 5, 8, and 14, respectively). For each example, the top-left shows the T1-weighted imaging; the top-right shows the FLAIR imaging; the bottom-left shows the FDG-PET; and the bottom-right shows the post-operative T1-weighted imaging. The red arrows indicate the MRI-visible lesion. The lesion and resection cavity were manually segmented in volume space and projected to the surface. The top row of surface maps shows the epicentre (MRI-visible FCD) in blue and the resection cavity in red. The middle row shows the SUVR maps. We calculated asymmetry maps and thresholded at < −0.2 to obtain cluster maps as shown in the bottom row. Of note, the hypometabolic cluster in the left frontal lobe of Patient 5 was due to a previous resection that failed to stop their seizures.

### Statistics

We conducted a paired t-test to compare SUVR in the epicentre compared to the border. Statistical significance was set at p <0.05.

A logistic mixed effects model was used to assess the relationship between presence of pathology and specimen location. The binary outcome was presence of pathology (fFCD/DNO vs Neg) and the primary predictor was location (Epicentre vs Border). We included a random intercept for each subject to account for repeated measures within individuals. The model was fit using the brm() function in the brms package in R statistical software. The model used a Bernoulli likelihood with a logit link and a default prior was used.

To investigate the association between anatomical location and pathological classification, we fitted a Bayesian multinomial logistic mixed-effects model with pathology (fFCD, DNO, or Neg) as outcome variable, location (Epicentre or Border) as predictor, and subject as a random effect. Negative pathology was set as the reference to allow for comparisons of the log-odds of fFCD and DNO relative to negative pathology. This analysis was repeated excluding three patients who had intracranial EEG investigations to assess whether the association between specimen location and histopathological classification was robust to the exclusion of patients with more subtle pathology requiring invasive presurgical monitoring.

To assess whether unequal specimen contributions influenced our findings, we repeated the primary analysis separately excluding the three patients contributing the most epicentre specimens and the three contributing the most border specimens. Furthermore, we performed a leave-one-subject-out sensitivity analysis. The Bayesian logistic mixed-effects model was refitted 14 times, sequentially excluding all specimens from one subject while retaining the remaining 13 subjects. The posterior mean and 95% credible interval were extracted from each refitted model and compared with estimates from the full cohort.

Finally, we performed additional exploratory analyses incorporating quantitative imaging measures into the model, including regional SUVR (Epicentre or Border), volume (Epicentre or Border), and proportion of hypometabolism resected as predictors.

## Results

Boxplots of the volumes of the epicentre (median 1.7± 3.2 mL), the border regions (median 5.3 ± 20.2 mL), and the resection cavity (median 6.6 ± 22.5 mL) are displayed in **Figure 3A**. A boxplot of the proportion of the hypometabolism resected is shown in **Figure 3B**. On average, 65.5±28.2% of the resection-associated hypometabolism was resected. We also compared the average SUVR in the epicentre versus the border in the boxplots in **Figure 3C**. The mean SUVR in the epicentre was 1.84±0.38, while the mean SUVR in the border was 2.24±0.42. A paired t-test was conducted, showing significantly lower SUVR in the epicentre compared to the border (t(13) = −5.79, p <0.001).

**Figure 3.**
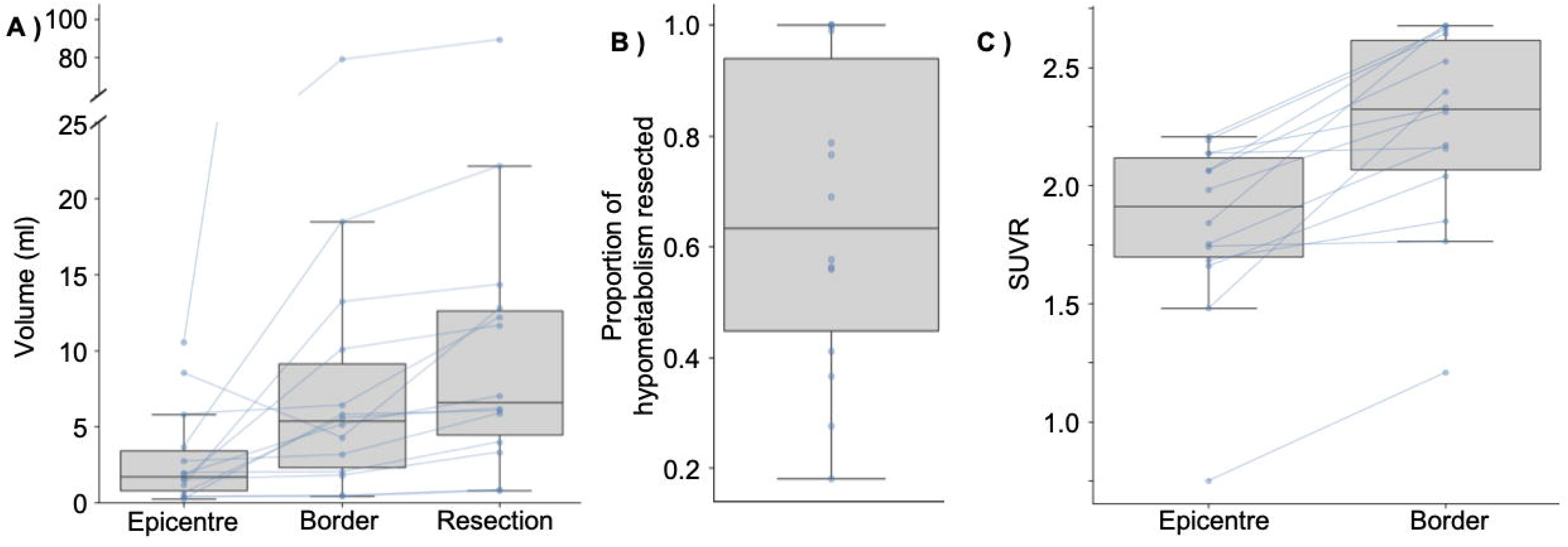
Boxplots summarizing imaging findings. The volume of the epicentre and border regions, and resection cavity are summarized in the boxplot in (A). Individual data points are plotted with lines connecting points of the same patient. The y-axis is broken to show the large border region and resection cavity for Patient 8. The distribution of proportion of hypometabolism resected is demonstrated in the boxplot in (B). The median proportion was 65.5% with a range of 18-100%. The distribution of SUVR for the epicentre and border are plotted in (C). Individual data points are plotted with lines connecting points of the same patient. Paired t-test demonstrated a significant difference in SUVR between the two regions (t(13) = −5.79, p <0.001). Boxplots show the median value in each subplot and boxplot whiskers extend to the maximum or minimum value within 1.5 times the interquartile range.

Intraoperatively, a total of 136 samples were taken from the epicentre of the lesion (N=64, median 4±3 specimens/patient, range 1-13) and the FDG-PET hypometabolic borders of the resection (N=72, median 5±4 specimens/patient, range 1-13).

All patients were found to have FCD II (no type I was seen in our series) (**Table 1**). Nine out of 14 had FCD IIa (64%) while 5/14 had FCD IIb (36%). There was on average 3.9±2.2 years of post-operative follow-up (all patients had at least 1-year follow-up; with 11/14 having >2-year follow-up) and all patients had Engel Ia status at the last known visit.

In the epicentre, 48/64 (75%) specimens had pathology (either fFCD or DNO), 38/64 (59%) had fFCD, 10/64 (16%) had DNO, and 16/64 (25%) were negative for pathology (**Figure 4**). In the hypometabolic borders, 44/72 (62%) specimens had pathology (either fFCD or DNO), 22/72 (31%) had fFCD, 22/72 (31%) had DNO, and 28/72 (38%) were negative for pathology. Of note, all patients had positive pathology in at least one of their epicentre specimens and in at least one of their hypometabolic border specimens (**Table 1**).

**Figure 4.**
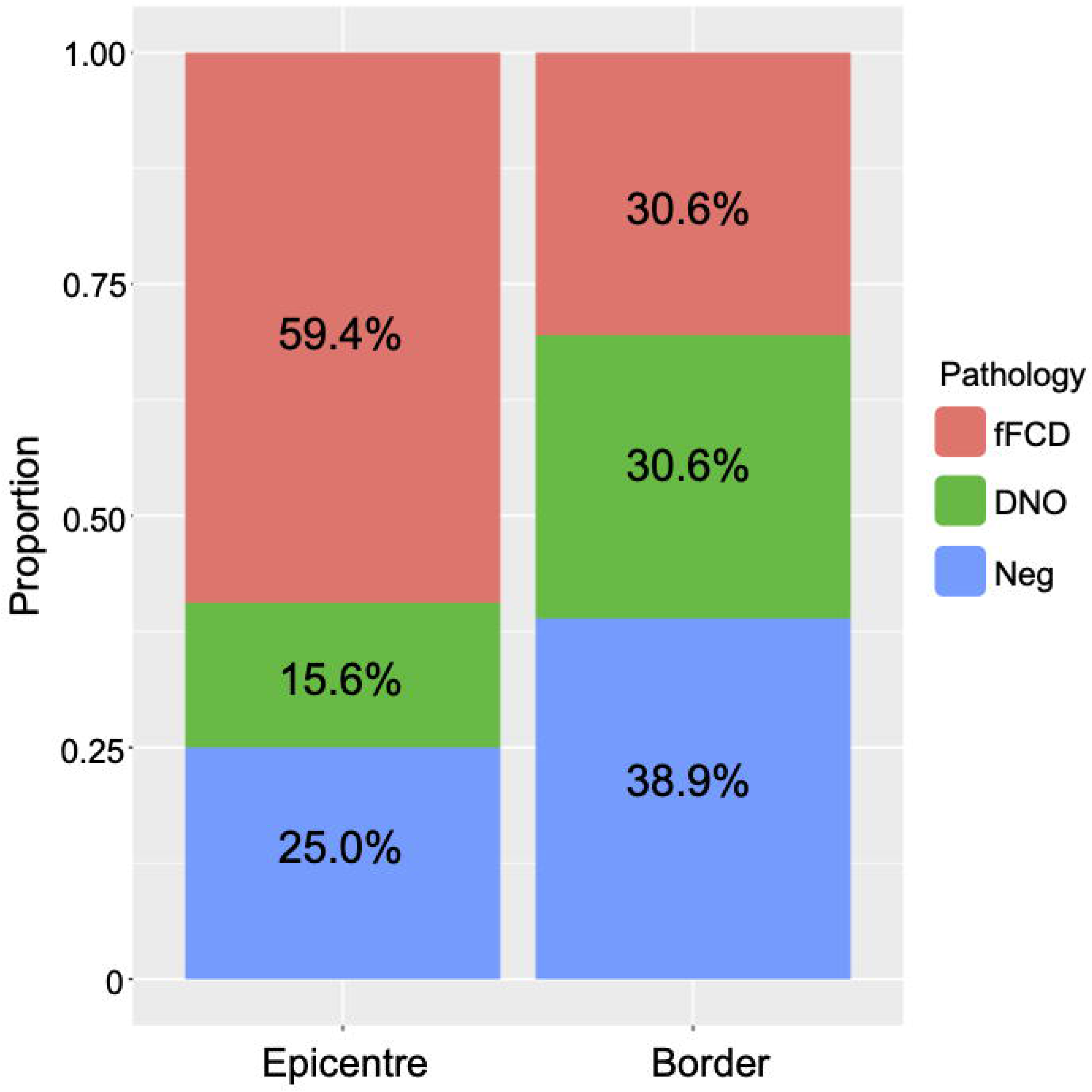
Bar plots showing the proportion and distribution of pathology in Epicentre and Border regions. The Epicentre had a majority FCD while Border regions had up to 62% pathological cells (either fFCD or DNO). DNO, dysmorphic neurons only; fFCD, frank focal cortical dysplasia; Neg, negative.

We fit a logistic mixed effects model with pathology as the dependent variable, location as the predictor, and subject as a random intercept. The model used for analysis was *Pathology ∼ location + (1|subject)*. The model was estimated using Bayesian sampling with 4 chains (2000 iterations each; 1000 warm up), yielding 4000 post-warmup draws.

The intercept was estimated at 1.30 (95% credible interval (CI): 0.60–2.10), indicating that in the epicentre, the odds of pathology were reliably greater than zero. In the border, pathology was less likely than in the epicentre (estimate −0.83, 95% CI: −1.72–0.00).

Next, we fit a Bayesian logistic mixed effects model to examine the relationship between pathology type (fFCD, DNO, negative) and specimen location (epicentre vs border), while accounting for inter-subject variability. The model used for analysis was *Pathology_type ∼ location + (1|subject)*. The model showed good convergence across parameters (all R = 1.00). Substantial between-subject variability was observed in baseline pathology probabilities (sd[μDN_Intercept] = 1.95, 95% CI [0.85, 3.78]; sd[μFCD_Intercept] = 0.43, 95% CI [0.02, 1.20]).

At the epicentre (reference location), the log odds of fFCD pathology were significantly higher relative to negative specimens (Estimate = 1.00, 95% CI [0.32, 1.77]), whereas the log odds of DNO pathology did not differ significantly from negative specimens (Estimate = –1.19, 95% CI [–2.91, 0.24]) (**Table 2**).

**Table 2.** Summary of Bayesian logistic mixed-effects model. Negative pathology was set as the reference to allow for comparisons of fFCD and DNO relative to negative pathology. CI, credible interval; OR, odds ratio. DNO, dysmorphic neurons only; fFCD, frank focal cortical dysplasia.

|  |  |  | 95% CI |  |
|---|---|---|---|---|
| Outcome logit | Parameter | Log OR | Lower | Upper |
| fFCD (vs Neg) | Intercept (Epicentre) | 1.00 | 0.32 | 1.77 |
|  | Border Location | -1.25 | -2.17 | -0.40 |
| DNO (vs Neg) | Intercept (Epicentre) | -1.19 | -2.9 | 0.24 |
|  | Border Location | 0.35 | -0.85 | 1.59 |

Location had a strong effect on the probability of fFCD pathology: specimens from the border regions were markedly less likely to show fFCD compared to the epicentre (Estimate = –1.25, 95% CI [–2.17, –0.40]). In contrast, the probability of DNO pathology did not significantly differ between border and epicentre specimens (Estimate = 0.35, 95% CI [–0.85, 1.59]).

Posterior predicted probabilities are illustrated in **Figure 5**. Specimens from the epicentre showed the highest estimated probability of fFCD pathology, while border specimens showed a reduced probability and a relatively higher probability of DNO pathology. These results support a spatial difference in distribution of histopathology, with fFCD histopathology concentrated in the center of the EZ and less abundant DNO extending toward the periphery.

**Figure 5.**
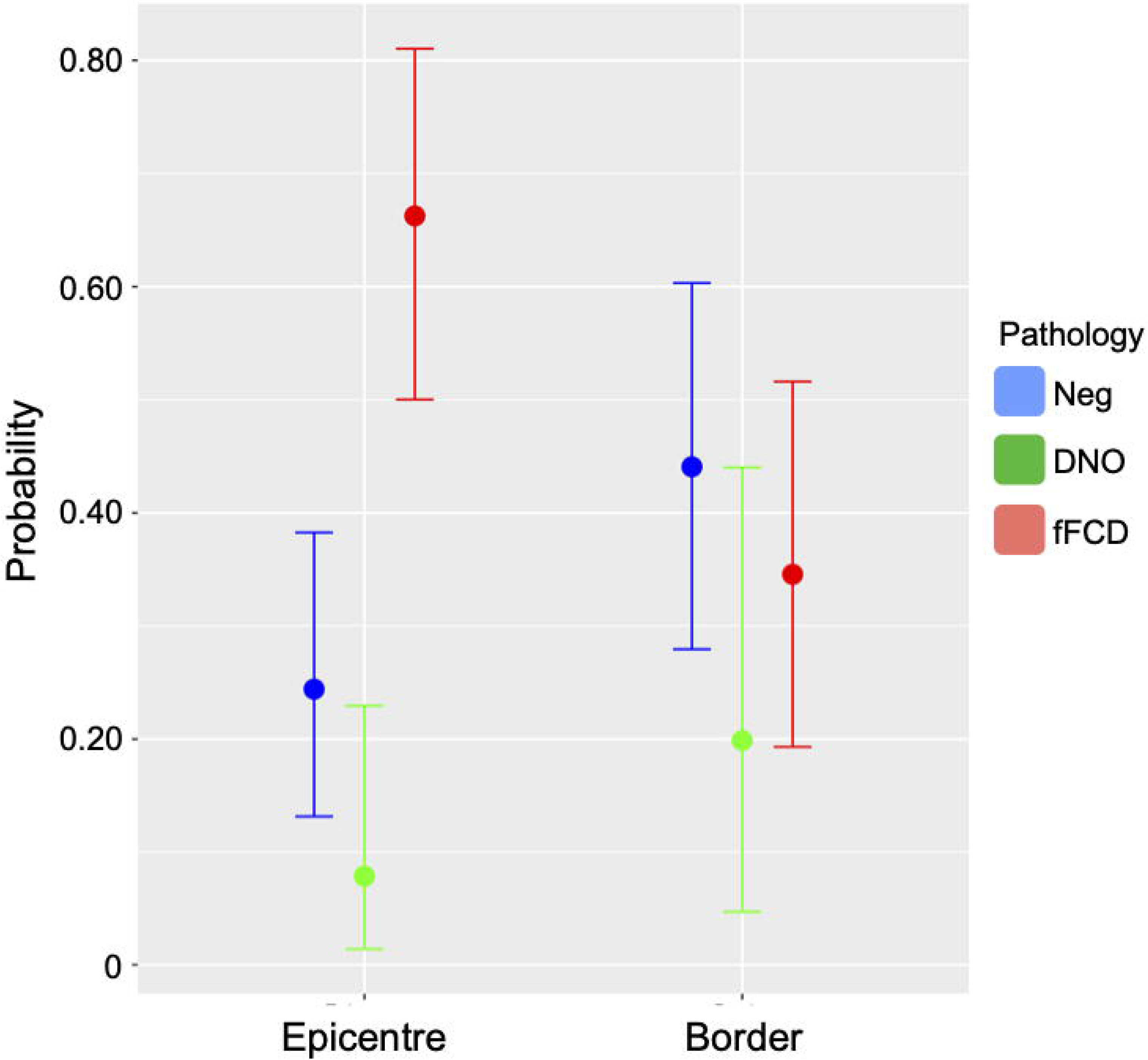
Posterior predicted probabilities from the Bayesian logistic mixed effects model. There was a high probability of fFCD in Epicentre regions that decreased in Border regions. There was a slightly higher chance of negative pathology in the Border region versus the Epicentre. DNO was roughly evenly probable in either location. DNO, dysmorphic neurons only; fFCD, frank focal cortical dysplasia; Neg, negative.

This analysis was repeated excluding three patients with intracranial EEG and we found that our main findings were robust to the exclusion of these patients (**Supplemental Table 1**).

To exclude the possibility that certain subjects were driving our effects, we repeated our main analysis separately excluding the top 3 patients contributing the most epicentre and border specimens. First, we excluded patients 9, 10, and 14, who collectively contributed 29/64 (45%) epicentre specimens. We found an association between location and odds of fFCD relative to negative pathology consistent with the original results and there was no clear evidence of a location effect for DNO relative to negative pathology (**Supplemental Table 2**). Next, we excluded patients 6, 9, and 13, who collectively contributed 31/72 (43%) border specimens. Again, our main findings were robust to the exclusion of these patients (**Supplemental Table 3**). Furthermore, we performed a leave-one-subject-out sensitivity analysis. Our analysis showed that the association between specimen location and histopathological findings was robust to the exclusion of individual subjects (**Figure 6**). The odds of fFCD relative to negative pathology in the epicentre (intercept) remained consistently above 0, while the odds of fFCD relative to negative pathology in the border remained consistently below 0. The 95% credible intervals excluded 0 in all iterations, indicating that the association was not driven by any one subject. In contrast the association between specimen location and DNO relative to negative pathology was less consistent. The exclusion of Patient 3 lead to the 95% credible interval to exclude 0. Otherwise, the 95% credible interval included 0 across all iterations, consistent with the full model.

**Figure 6.**
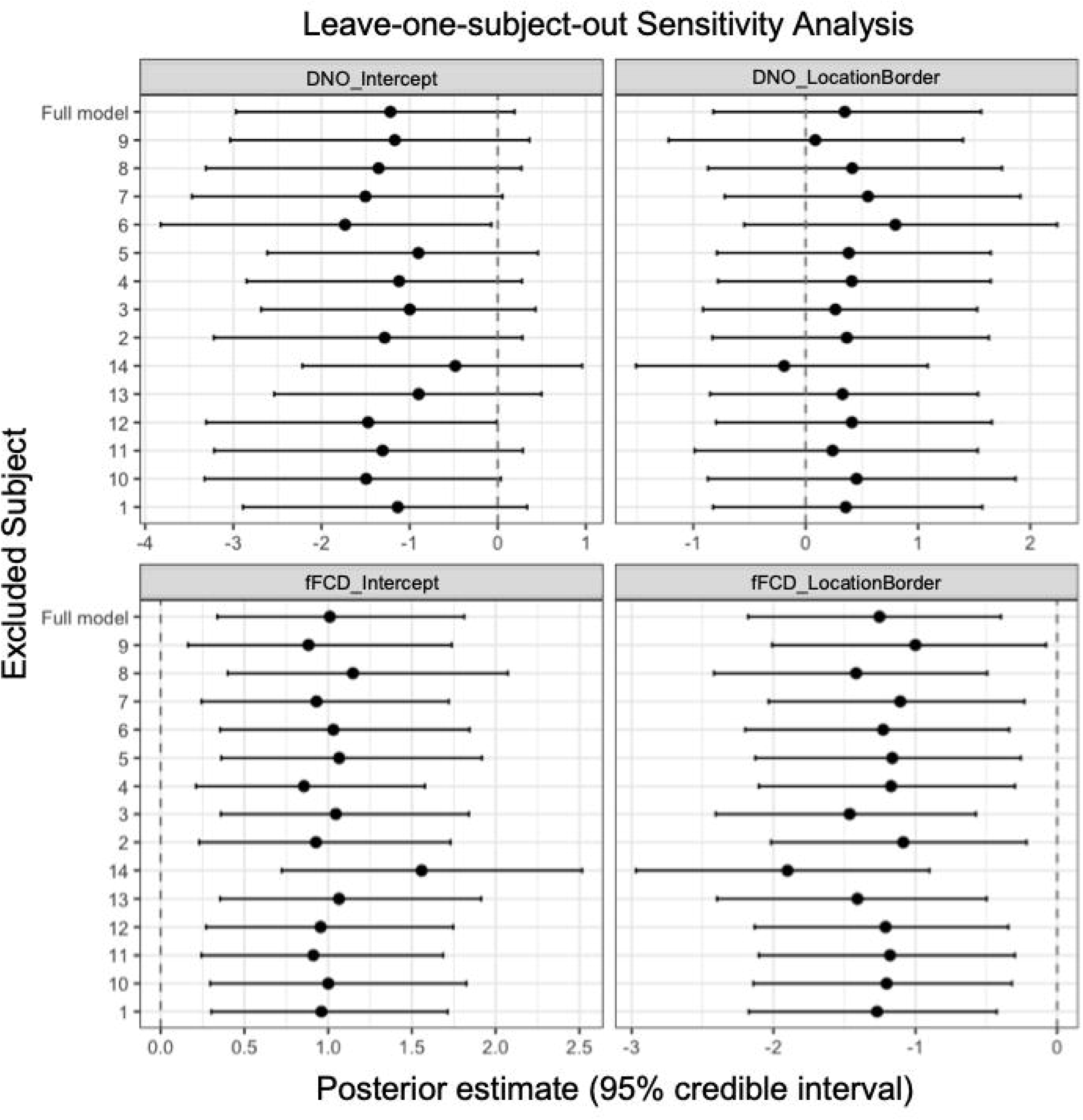
Leave-one-subject-out sensitivity analysis. In order to establish that the effects seen in the main analysis were not driven by any single subject, we reran the model iteratively, excluding one patient each run. The 95% credible interval included 0 for all but one iteration for DNO and excluded 0 for all iterations for fFCD. These results were consistent with the full model, indicating our results were relatively robust to the exclusion of individual patients.

To test the association between histopathology, location, and imaging markers, we performed an additional exploratory analysis including Z-scored regional SUVR, volume, and proportion of hypometabolism in the model. The model used for analysis was *Pathology_type ∼ location + SUVR_z + volume_z + proportion_hypometabolism_resected_z + (1|subject)*. The model showed good convergence (all parameter R = 1.00). Substantial between-subject variability was observed in baseline pathology probabilities (sd[μDN_Intercept] = 1.96, 95% CI [0.68, 4.18]; sd[μFCD_Intercept] = 0.47, 95% CI [0.01, 1.16]).

The effects seen in the main analysis (**Table 2**) were maintained (**Table 3**). Neither standardized regional SUVR nor volume had a significant effect on pathology. Standardized proportion of hypometabolism resected showed a statistically credible inverse association with fFCD status (Estimate = −0.72, 95% CI [-1.39, −0.07]). No association was seen with DNO status (Estimate = −0.49, 95% CI [-2.39, 1.07]).

**Table 3.** Summary of exploratory Bayesian logistic mixed effects model incorporating imaging findings. Negative pathology was set as the reference to allow for comparisons of fFCD and DNO relative to negative pathology. Regional SUVR and volume, and proportion of hypometabolism resected were standardized prior to inclusion in the model. CI, credible interval; OR, odds ratio. DNO, dysmorphic neurons only; fFCD, frank focal cortical dysplasia; SUVR, standardized uptake value ratio.

|  |  |  | 95% CI |  |
|---|---|---|---|---|
| Outcome logit | Parameter | Log OR | Lower | Upper |
| fFCD (vs Neg) | Intercept (Epicentre) | 1.42 | 0.54 | 2.41 |
|  | Border Location | -2.06 | -3.65 | -0.62 |
|  | SUVR | 0.39 | -0.41 | 1.19 |
|  | Volume | 0.12 | -0.39 | 0.65 |
|  | Proportion of hypometabolism resected | -0.72 | -1.39 | -0.07 |
| DNO (vs Neg) | Intercept (Epicentre) | -1.49 | -3.44 | 0.12 |
|  | Border Location | 0.70 | -1.33 | 2.82 |
|  | SUVR | -0.32 | -1.61 | 1.22 |
|  | Volume | -0.25 | -1.07 | 0.51 |
|  | Proportion of hypometabolism resected | -0.49 | -2.39 | 1.07 |

## Discussion

In this study of pediatric focal epilepsy patients who underwent resective surgery for FCD, we provide evidence for the presence of potentially epileptogenic cells in the peripheral area of hypometabolism outside of the MRI-defined lesion. Epicentre regions with MRI alterations were more likely to have fFCD than negative pathology, while border regions were less likely to have fFCD. Areas of less abundant DNO were present in both epicentre and border regions without strong evidence of a location-based difference. Overall, this suggests a spatial difference in distribution of histopathology throughout the area of hypometabolism with a significant proportion of border specimens also containing pathology (up to 62%).

This spatial difference in distribution of histopathology may have associations with the degree of hypometabolism and electro-clinical features. In patients with malformations of cortical development, Lagarde and colleagues examined FDG-PET in lesional, epileptogenic non-lesional, propagation, and non-involved brain regions as defined by stereotactic EEG.^24^ They demonstrated a gradient of worsening FDG-PET hypometabolism, from non-involved areas having the highest metabolism to lesional areas having the lowest metabolism (i.e., most hypometabolic). Whether this electroclinical/FDG-PET gradient is underlain by a gradient of pathology remains to be elucidated with further quantitative analyses.

In our exploratory analysis, despite finding a significant difference in hypometabolism between the epicentre and the border, we were unable to demonstrate a clear association between pathology and regional SUVR in our modeling. Because the precise specimen locations could not be resolved retrospectively, specimen specific SUVR could not be obtained and we had to incorporate the regional (epicentre/border) SUVR in our model. No association was seen between epicentre/border volume or pathology either, which may reflect the distinction between the overall extent of the imaging abnormality and its microscopic composition. On the other hand, we did observe a credible inverse association between proportion of hypometabolism resected and fFCD status. More complete coverage of the hypometabolic region could capture a broader mixture of tissue, including cortex without frank FCD. Because fFCD was concentrated in the epicentre, specimens from patients with a greater proportion of hypometabolism resected may contain relatively more negative tissue.

Our group’s previous work has shown a lack of association between histopathology and presence of interictal epileptic discharges, ripples, and fast ripples on stereotactic EEG, implying that the EZ may not be strictly a histopathological entity, but rather a hybrid of abnormal cells and normal-appearing cells.^25^ Similarly, in the present work, we show that the area of hypometabolism beyond the lesion epicentre contains a combination of normal and dysplastic cells. It is possible that the area of hypometabolism delineates the same epileptogenic hybrid area as high frequency oscillations (HFOs). Zhao and colleagues found a significant relationship between degree of hypometabolism and HFO generation rate, suggesting a common pathophysiologic mechanism.^26^ This association could help guide SEEG insertion planning and provide prognostic information on post-operative seizure and functional outcome. Another study examining the localizing value of FDG-PET in patients operated for FCD showed improved sensitivity for EZ localization when combining PET/MRI and electroclinical data.^27^ The typical electrical signature of FCD type 2 was found in the area of greatest hypometabolism in each patient and the authors found in their series that invasive monitoring could be avoided in some cases where structural imaging was ambiguous. Beyond its predictive value for post-surgical outcomes, FDG-PET abnormalities may also be linked to functional outcomes as well. In a study of 15 patients who underwent surgical resection for FCD in the Rolandic area, FDG-PET metabolism was positively associated with postoperative motor deficits (i.e., resecting areas of pre-existing hypometabolism did not cause such deficits),^28^ suggesting that hypometabolism reflects functional reorganization in primary motor cortex.

Seizure-freedom among our cohort of 14 patients was higher (100% Engel Ia at 1 year post-surgery) than that typically reported for FCD.^4^ Several factors likely contributed to the favorable surgical outcomes observed in our cohort. First, all patients had a visible FCD identified upon re-review of imaging, and resections did not involve eloquent cortical areas. Second, the relatively short follow-up period for some patients (e.g., one year) may have inflated the apparent success rate; it remains to be seen whether these patients develop recurrent seizures later in their post-operative course. Finally, rather than defining the surgical margins solely by the MRI-visible lesion, we targeted the broader area of PET-defined hypometabolism as the basis for the resection. Previous studies have shown that dysmorphic neurons in fact contribute to interictal spikes, fast gamma activity, and ripples.^29^ While frank FCD may be found in the borders of the hypometabolic areas, ostensibly even just the presence of dysmorphic neurons may be enough to contribute to epileptogenicity. By maximizing the extent of hypometabolism in the surgical resection, we are likely removing more potentially epileptogenic cells and thus improving the chances of successful surgical outcome.

Previous work on non-lesional temporal lobe epilepsy has shown that extent of resection of the hypometabolic tissue is associated with improved post-surgical outcome.^30^ In extra-temporal lobe epilepsy, there is conflicting data on extent of resection of hypometabolic abnormality and surgical outcomes. One study by Juhasz and colleagues on pediatric patients with drug resistant neocortical epilepsy showed no correlation between extent of non-resected hypometabolism and post-operative seizure freedom.^31^ Another study by Zhao and colleagues demonstrated that the extent of resection of hypometabolism did not differ between seizure-free and non-seizure-free patients.^26^ However, resecting tissue that had both fast ripples and hypometabolism was significantly predictive. On the other hand, Sainburg and colleagues have shown that surgical targeting of lateralized FDG-PET hypometabolism is associated with long term post-surgical outcomes.^32^ This effect was seen in TLE patients, but also generalized to a subcohort of extra-temporal lobe epilepsy patients. More recently, Gijs et al., using an anatomy-corrected asymmetry index, demonstrated that patients with Engel I outcome had a higher percentage of their hypometabolic volume resected independent of MRI lesion status or FCD subtype.^33^ Beyond extent of resection, it appears that the pattern of hypometabolism is also an important predictor of outcome. In a cohort of neocortical epilepsy patients, Wong et al. showed that a larger volume of remote hypometabolism (i.e. more widespread abnormality) was a predictor of surgical failure.^34^ This was reinforced in a recent meta-analysis that suggested that diffuse hypometabolism was associated with worse surgical outcomes compared to focal hypometabolism. Overall, further work is required to better understand the spatial distribution and burden of pathology within the hypometabolic abnormality and its relationship with surgical outcomes.

We had three patients who had subtle imaging findings who required intracranial recordings: two had SEEG insertions while one had subdural grids and depth electrodes. Following detection of the seizure onset zone, a subtle lesion could be retrospectively identified on imaging. Among these three patients, the epicentre specimens showed fFCD in 4/4 specimens from patient 4 and 3/5 specimens from patient 5, whereas both epicentre specimens from patient 6 showed DNO. Thus, invasive monitoring was not uniformly associated with more subtle specimen-level pathology, although the small subgroup precludes firm conclusions. The overarching resection strategy was consistent across patients. We aimed to resect the MRI-defined lesion and surrounding FDG-PET hypometabolic tissue when safely feasible. Invasive EEG provided additional localization information in these three cases. Repeating our analysis excluding these patients yielded consistent results, indicating our main findings were not driven by patients with subtle imaging findings.

Our study has a number of limitations. First, our sample size was modest with only 14 patients. However, we had a total of 136 specimens across the cohort with an average of 9.7 specimens per patient. Our sample sizes were unbalanced across subjects with 45% of epicentre specimens coming from a group of 3 patients and 43% of border specimens coming from another group of 3 patients. Our sensitivity analyses excluding these patients show similar results to our main findings. In addition, with our leave-one-subject-out analysis, we show that our findings were robust to the exclusion of single subjects. Overall, these findings indicate that our main results are likely not driven by any one subject or groups of subjects with disproportionate number of samples. Another limitation was that our pathological sampling was restricted to the extent of resection. Within the limitations of current day research tools in patients, it is impossible to know if pathological cells may be present beyond the area of resection, and if so, in what proportion in comparison to the resected tissue. In keeping with the spatial difference in distribution of histopathology seen in our study, it may be reasonable to infer that few pathological cells may present at the edge of the resection cavity, and certainly too few to cross the threshold of epileptogenicity given the seizure-free status of our cohort. The qualitative determination of pathology and FDG-PET hypometabolism (at time of surgery) is another limitation. Pathology was based on descriptions of the neuropathologists’ reports and quantitative measures of FCD and dysmorphic neurons were not obtained. Similarly, the area of hypometabolism on which the resection was planned was delineated manually. This was done in a consistent manner for all cases by one person (the lead surgeon) using only the MRI/FDG-PET co-registered images. However, this remains subjective and potentially biased by adjacent MRI signal abnormalities, anatomical borders, and quality of the FDG-PET images, as well as other prior clinical and/or electrophysiological knowledge regarding the case (e.g., semiology, MEG results, and EEG/SEEG findings). Finally, we only had patients with an MRI-visible lesion and FCD type II histopathology in our cohort. Whether our findings are generalizable to those with an MRI-negative lesion and/or FCD type I histopathology remains to be seen.

Future studies will continue further prospective data collection and compare pathology between seizure-free and non-seizure free patients to assess the effect of resection border pathology on post-surgical outcomes. A study on examining genetic data on removed stereotactic EEG electrodes is ongoing and may highlight associations between pathogenic variant load and metabolism in the EZ and surrounding areas.^35^ Future studies could also incorporate more precise localization of specimen sampling to enable direct correlation of pathological findings with structural/functional MRI and PET findings. Finally, quantification of pathology will enable assessment of potential associations between amount of pathological cell burden with metabolism.

Overall, in this study of pediatric focal epilepsy patients whose resection borders were informed largely by the preoperative FDG-PET, we show that pathological cells (either frank FCD or dysmorphic neurons) are present in these hypometabolic borders beyond the MRI-detected signal abnormalities. This suggests that FDG-PET may be useful in delineating the histopathological borders of FCD, which we know extend beyond what is seen on MRI. Thus, maximal, safe resection of the area of hypometabolism may remove these potentially epileptogenic cells, thus improving post-operative surgical outcome.

## Acknowledgements

J.L. is funded by the Fonds de la Recherche du Québec – Santé (FRQS) and the Ministère de la Santé et des Services sociaux du Québec (MSSS). R.W.R.D. is funded by the Fonds de la Recherche du Québec – Santé (FRQS) Chercheur Boursiers Clinicien Program. B.C.B. acknowledges support from the Canadian Institutes of Health Research, CIHR (FDN-154298, PJT-174995, PJT-206196, CIHR PJT-203761, CIHR PJT-191853), SickKids Foundation (NI17-039), Natural Sciences and Engineering Research Council (NSERC RGPIN-2025-05932), Azrieli Center for Autism Research of the Montreal Neurological Institute (ACAR), BrainCanada, FRQ-S, the Helmholtz International BigBrain Analytics and Learning Laboratory (Hiball), Healthy Brains and Healthy Lives (HBHL), Centre for Aging + Brain Health Innovation (CABHI), the Canada Research Chairs Program (CRC), and the Centre of Excellence in Epilepsy at the Neuro (CEEN).

## Author Contributions

Study design and conception: R.W.R.D, J.L.; data acquisition, analysis, interpretation: R.W.R.D, J.L., N.v.E., M.H., Y.R., M.-C.G., J.K., B.C.B.; manuscript drafting: R.W.R.D, J.L. All authors provided feedback and approved of the final manuscript.

## Ethical Publication Statement

B.C.B. is a co-founder of BrainScores Inc. We confirm that we have read the Journal’s position on issues involved in ethical publication and affirm that this report is consistent with those guidelines.

---

**Supplemental Figure 1.** Surface maps showing the MRI-visible FCD, resection cavity, and hypometabolic clusters for each patient. For each patient, the top row shows the epicentre in blue and the resection cavity in red, while the bottom row shows the hypometabolic cluster map in purple (thresholded at asymmetry index <-0.2).

**Supplemental Table 1.** Summary of Bayesian logistic mixed-effects model examining the association between pathology type and specimen location, while accounting for inter-subject variability and excluding three patients who had intracranial electro-encephalography investigations. CI, credible interval; OR, odds ratio. DNO, dysmorphic neurons only; fFCD, frank focal cortical dysplasia.

|  |  |  | 95% CI |  |
|---|---|---|---|---|
| Outcome logit | Parameter | Log OR | Lower | Upper |
| fFCD (vs Neg) | Intercept (Epicentre) | 0.92 | 0.19 | 1.71 |
|  | Border | -1.00 | -2.06 | -0.03 |
| DNO (vs Neg) | Intercept (Epicentre) | 0.92 | -3.1 | 0.19 |
|  | Border | 0.89 | -0.49 | 2.35 |

**Supplemental Table 2.** Summary of Bayesian logistic mixed-effects model examining the association between pathology type and specimen location, while accounting for inter-subject variability and excluding the top three patients (9, 10, 14) with the most epicentre specimens (29/64 (45%)). CI, credible interval; OR, odds ratio. DNO, dysmorphic neurons only; fFCD, frank focal cortical dysplasia.

|  |  |  | 95% CI |  |
|---|---|---|---|---|
| Outcome logit | Parameter | Log OR | Lower | Upper |
| fFCD (vs Neg) | Intercept (Epicentre) | 1.51 | 0.51 | 2.64 |
|  | Border | -1.68 | -2.83 | -0.56 |
| DNO (vs Neg) | Intercept (Epicentre) | -0.59 | -2.55 | 1.12 |
|  | Border | -0.53 | -2.09 | 1.06 |

**Supplemental Table 3.**
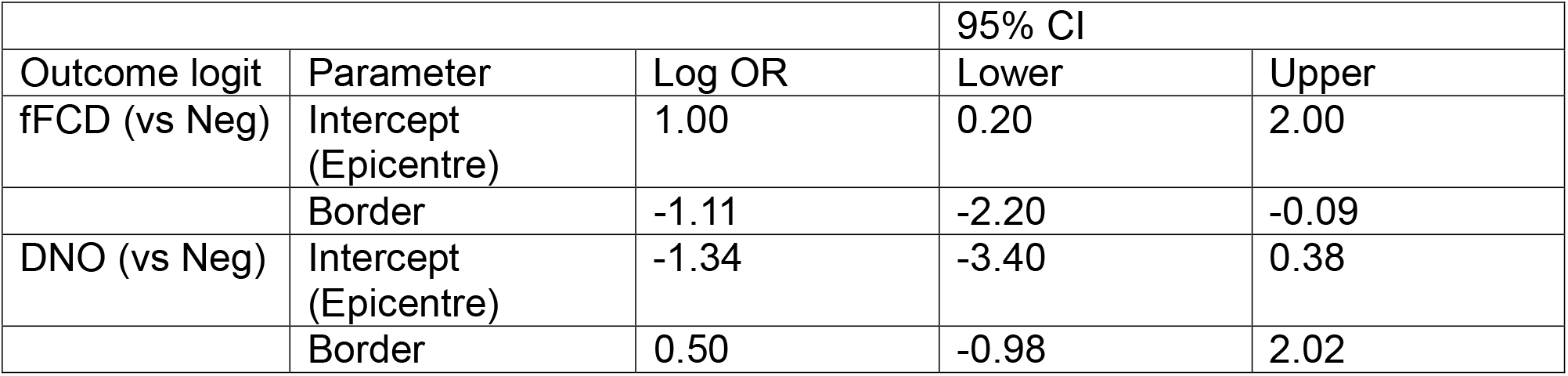
Summary of Bayesian logistic mixed-effects model examining the association between pathology type and specimen location, while accounting for inter-subject variability and excluding the top three patients (6, 9, 13) with the most border specimens (31/72 (43%)). CI, credible interval; OR, odds ratio. DNO, dysmorphic neurons only; fFCD, frank focal cortical dysplasia.

|  |  |  | 95% CI |  |
|---|---|---|---|---|
| Outcome logit | Parameter | Log OR | Lower | Upper |
| fFCD (vs Neg) | Intercept (Epicentre) | 1.00 | 0.20 | 2.00 |
|  | Border | -1.11 | -2.20 | -0.09 |
| DNO (vs Neg) | Intercept (Epicentre) | -1.34 | -3.40 | 0.38 |
|  | Border | 0.50 | -0.98 | 2.02 |

## Notes

### Summary of Updates

Additional analyses and figures/tables; expanded discussion

## References

1. Lopez-Rivera, J. A. et al. Incidence and prevalence of major epilepsy-associated brain lesions. Epilepsy Behav. Rep. 18, 100527 (2022).

2. Najm, I. et al. The ILAE consensus classification of focal cortical dysplasia: An update proposed by an ad hoc task force of the ILAE diagnostic methods commission. Epilepsia 63, 1899–1919 (2022).

3. Blümcke, I. et al. The clinicopathologic spectrum of focal cortical dysplasias: A consensus classification proposed by an ad hoc Task Force of the ILAE Diagnostic Methods Commission. Epilepsia 52, 158–174 (2011).

4. Martinez-Lizana, E. et al. Long-term seizure outcome in pediatric patients with focal cortical dysplasia undergoing tailored and standard surgical resections. Seizure 62, 66–73 (2018).

5. Sisodiya, S. M., Fauser, S., Cross, J. H. & Thom, M. Focal cortical dysplasia type II: biological features and clinical perspectives. Lancet Neurol. 8, 830–843 (2009).

6. Rubí, S. et al. Validation of FDG-PET/MRI coregistration in nonlesional refractory childhood epilepsy. Epilepsia 52, 2216–2224 (2011).

7. Salamon, N. et al. FDG-PET/MRI coregistration improves detection of cortical dysplasia in patients with epilepsy. Neurology 71, 1594–1601 (2008).

8. Lerner, J. T. et al. Assessment and surgical outcomes for mild type I and severe type II cortical dysplasia: a critical review and the UCLA experience. Epilepsia 50, 1310– 1335 (2009).

9. Ollenberger, G. P. et al. Assessment of the role of FDG PET in the diagnosis and management of children with refractory epilepsy. Eur. J. Nucl. Med. Mol. Imaging 32, 1311–1316 (2005).

10. Courtney, M. R. et al. Association of Localizing 18F-FDG-PET Hypometabolism and Outcome Following Epilepsy Surgery. Neurology 102, e209304 (2024).

11. da Silva, E. A., Chugani, D. C., Muzik, O. & Chugani, H. T. Identification of frontal lobe epileptic foci in children using positron emission tomography. Epilepsia 38, 1198–1208 (1997).

12. Gaillard, W. D. et al. FDG-PET in children and adolescents with partial seizures: Role in epilepsy surgery,evaluation. Epilepsy Res. 20, 77–84 (1995).

13. Semah, F. et al. Is interictal temporal hypometabolism related to mesial temporal sclerosis? A positron emission tomography/magnetic resonance imaging confrontation. Epilepsia 36, 447–456 (1995).

14. Rho, J. M. & Boison, D. The metabolic basis of epilepsy. Nat. Rev. Neurol. 18, 333– 347 (2022).

15. Liotta, A., et al. Metabolic Adaptation in Epilepsy: From Acute Response to Chronic Impairment. Int. J. Mol. Sci. 25, 9640 (2024).

16. Tenney, J. R., Rozhkov, L., Horn, P., Miles, L. & Miles, M. V. Cerebral glucose hypometabolism is associated with mitochondrial dysfunction in patients with intractable epilepsy and cortical dysplasia. Epilepsia 55, 1415–1422 (2014).

17. Schur, S. et al. New interinstitutional, multimodal presurgical evaluation protocol associated with improved seizure freedom for poorly defined cases of focal epilepsy in children. J. Neurosurg. Pediatr. 29, 74–82 (2022).

18. Goodall-Halliwell, I. et al. MICAFlow: Fast and robust MRI preprocessing bridging research neuroimaging and clinical practice. Imaging Neurosci. 4, IMAG.a.1366 (2026).

19. Seo, J. et al. Automated segmentation of postsurgical resection cavities on magnetic resonance imaging in focal epilepsy: A Multicentre Epilepsy Lesion Detection study. Epilepsia epi.70450 (2026) doi:10.1002/epi.70450.

20. Henschel, L. et al. FastSurfer - A fast and accurate deep learning based neuroimaging pipeline. NeuroImage 219, 117012 (2020).

21. Henschel, L., Kügler, D. & Reuter, M. FastSurferVINN: Building resolution-independence into deep learning segmentation methods-A solution for HighRes brain MRI. NeuroImage 251, 118933 (2022).

22. Thomas, B. A. et al. PETPVC: a toolbox for performing partial volume correction techniques in positron emission tomography. Phys. Med. Biol. 61, 7975–7993 (2016).

23. Marcus, D. S. et al. Informatics and Data Mining Tools and Strategies for the Human Connectome Project. Front. Neuroinformatics 5, (2011).

24. Lagarde, S. et al. Relationship between PET metabolism and SEEG epileptogenicity in focal lesional epilepsy. Eur. J. Nucl. Med. Mol. Imaging 47, 3130–3142 (2020).

25. von Ellenrieder, N. et al. No association between histopathology and neurophysiology in surgical specimens from pediatric focal epilepsy patients. Epilepsia 66, 3168–3179 (2025).

26. Zhao, B. et al. Interictal HFO and FDG-PET correlation predicts surgical outcome following SEEG. Epilepsia 64, 667–677 (2023).

27. Desarnaud, S. et al. 18F-FDG PET in drug-resistant epilepsy due to focal cortical dysplasia type 2: additional value of electroclinical data and coregistration with MRI. Eur. J. Nucl. Med. Mol. Imaging 45, 1449–1460 (2018).

28. Luo, W. et al. Predicting postoperative motor outcomes in the surgical management of Rolandic focal cortical dysplasia: the role of glucose metabolism. BMC Med. 23, 390 (2025).

29. Rampp, S. et al. Dysmorphic neurons as cellular source for phase-amplitude coupling in Focal Cortical Dysplasia Type II. Clin. Neurophysiol. 132, 782–792 (2021).

30. Vinton, A. B. et al. The extent of resection of FDG-PET hypometabolism relates to outcome of temporal lobectomy. Brain 130, 548–560 (2007).

31. Juhász, C. et al. Relationship of flumazenil and glucose PET abnormalities to neocortical epilepsy surgery outcome. Neurology 56, 1650–1658 (2001).

32. Sainburg, L. E. et al. Surgical targeting of lateralized 18F-fluorodeoxyglucose positron emission tomography hypometabolism relates to long-term epilepsy surgery outcomes. Epilepsia 66, 2816–2829 (2025).

33. Gijs, J. et al. Anatomy-corrected metabolic asymmetry predicts seizure freedom after surgery in focal cortical dysplasia. Eur. J. Nucl. Med. Mol. Imaging 53, 5088–5100 (2026).

34. Wong, C. H. et al. Relationship between preoperative hypometabolism and surgical outcome in neocortical epilepsy surgery. Epilepsia 53, 1333–1340 (2012).

35. Krochmalnek, E. et al. mTOR Pathway Somatic Pathogenic Variants in Focal Malformations of Cortical Development: Novel Variants, Topographic Mapping, and Clinical Outcomes. Neurol. Genet. 9, e200103 (2023).

