## Supplementary figures and images for "Pathology in resected areas of [^18^F]FDG PET hypometabolism in pediatric epilepsy patients with focal cortical dysplasia type II"

### Supplemental Figure 1

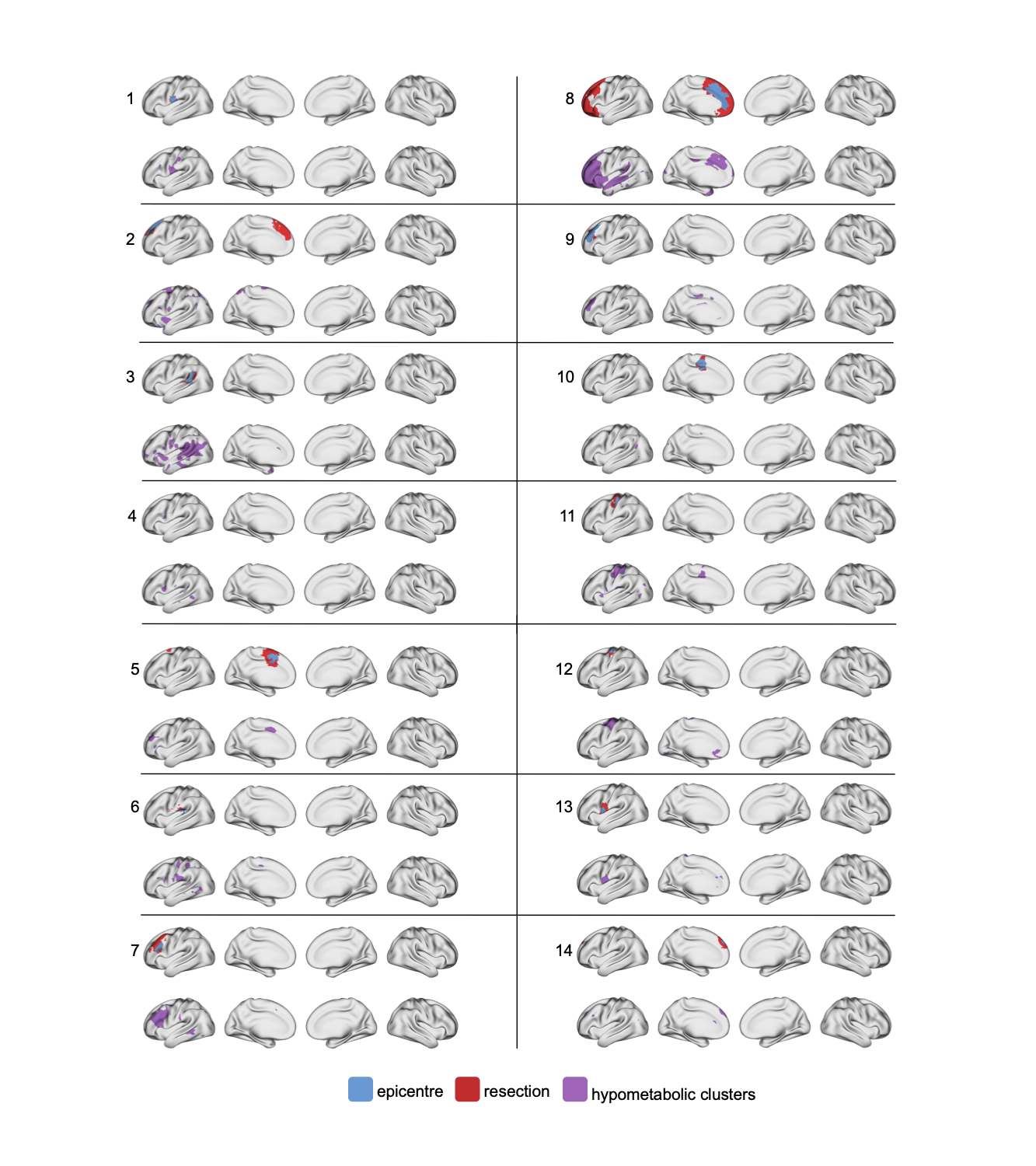
